# Leveraging genetic data to investigate molecular targets and drug repurposing candidates for treating alcohol use disorder and hepatotoxicity

**DOI:** 10.1101/791723

**Authors:** Joshua C. Gray, Mikela Murphy, Lorenzo Leggio

## Abstract

**Background:** Novel treatments for alcohol use disorder (AUD) and alcohol-related liver disease (ALD) are greatly needed. Genetic information can improve drug discovery rates by facilitating the identification of novel biological targets and potential drugs for repurposing.

**Methods:** The present study utilized a recently developed Bayesian approach, Integrative Risk Gene Selector (iRIGS), to identify additional risk genes for alcohol consumption using SNPs from the largest alcohol consumption GWAS to date (*N* = 941,280). iRIGS incorporates several genomic features and closeness of these genes in network space to compute a posterior probability for protein coding genes near each SNP. We subsequently used the Target Central Resource Database to search for drug-protein interactions for these newly identified genes and previously identified risk genes for alcohol consumption.

**Results:** We identified several genes that are novel contributions to the previously published alcohol consumption GWAS. Namely, *ACVR2A*, which is critical for liver function and linked to anxiety and cocaine self-administration, and *PRKCE*, which has been linked to alcohol self-administration. Notably, only a minority of the SNPs (18.4%) were linked to genes with confidence (≥.75), underscoring the need to apply multiple methods to assign function to loci. Finally, several previously identified risk genes for alcohol consumption code for proteins that are implicated in liver function and are targeted by drugs that are promising candidates for managing hepatotoxicity (e.g., metformin).

**Conclusions:** This study demonstrates the value of incorporating regulatory information and drug-protein interaction data to highlight additional molecular targets and drug repurposing candidates for treating AUD and ALD.

## 1. Introduction

Improving the treatment of patients with alcohol use disorder (AUD) and alcohol-related liver disease (ALD) is of vital importance from a clinical and public health standpoint (Leggio and Lee, 2017). Only three medications have been approved by the FDA to treat AUD – the last approval took place almost 15 years ago. No medications are approved for ALD. Targeting disease mechanisms with genetic support can increase success in drug development (Nelson et al., 2015). However, translating genome-wide association studies (GWASs) of complex diseases to target discovery and medication development remains challenging (Oprea et al., 2018). AUD is exemplary of this challenge; numerous large GWASs have yielded many significant SNPs (Kranzler et al., 2019; Liu et al., 2019), yet limited drug targets for treating AUD and alcohol-related consequences, such as ALD, have been identified.

We conducted two sets of analyses to identify additional risk genes for alcohol consumption and assess target druggability. First, we applied a modified version of the Integrative Risk Gene Selector (iRIGS) (Wang et al., 2019) to 98 genome-wide significant SNPs from the largest alcohol consumption GWAS to date, which assessed drinks per week (*M* = 7.84) in 941,280 individuals (Liu et al., 2019). In this study, iRIGS ranked genes at each SNP by integrating evidence from distal regulatory elements-promoter links to yield potential risk genes for alcohol consumption. Second, we classified gene encoded protein druggability based on the Target Development/Druggability Level (TDL) classification system (Oprea et al., 2018) from the Target Central Resource Database (TCRD).

## 2. Material and methods

### 2.1. Integrative Risk Gene Selector (iRIGS)

For the first set of analyses, we applied a modified version of iRIGS to 98 genome-wide significant SNPs (*p* < 5 × 10^−8^) associated with drinks per week from the GSCAN study (Liu et al., 2019) (rs7074871 was excluded because it had no nearby protein-coding genes). Briefly, iRIGS integrates two layers of information: 1) genomic features (i.e., distance from gene to SNP and four sets of regulatory connections derived from distal regulatory elements-promoter links from Hi-C, capture Hi-C, and FANTOM5 data) for genes within a 1 megabase (Mb) flanking region of each SNP; and 2) closeness of each gene in the network (Wang et al., 2019). Our modified version of the iRIGS method did not include de novo mutation enrichment or differential expression because the datasets used for the latter were specific to schizophrenia, there are no equivalent datasets to our knowledge for alcohol consumption, and there are many non-brain tissues relevant to alcohol consumption. We defined the cutoff for potential risk genes to be a posterior probability of ≥.75, indicating the gene is 75% likely to be related to the SNP (the probabilities of all genes within ±1Mb of a given SNP add up to 100%).

### 2.2. Target Development/Druggability Level (TDL)

For the second set of analyses, we incorporated genes identified in the iRIGS analyses and the analyses conducted in the prior GWAS. The methods for generating these alcohol consumption genes are discussed in detail in the prior study (Liu et al., 2019). Briefly, they defined a gene as implicated if it harbored variation of LD *r*^2^ > .3 with a genome-wide significant SNP or if it was located within 500kb of the SNP and was significant by the PASCAL gene-based test. This yielded 307 unique genes for our analyses. In order to identify the overlap of new and previously identified risk genes for alcohol consumption with preexisting drugs, we integrated drug-protein interaction information from the TCRD (Oprea et al., 2018). The TCRD defines TDLs according to 4 levels of confidence: T_clin_, targets have approved drug(s) with known mechanism(s) of action; T_chem_ targets have drugs or small molecules that satisfy activity thresholds; T_bio_ targets have no known drugs or small molecules that satisfy thresholds, but have Gene Ontology (GO) leaf term annotations, Online Mendelian Inheritance in Man (OMIM) phenotypes, or meet two of the three conditions: a fractional PubMed count > 5, > 3 National Center for Biotechnology Gene Reference Intro Function annotations, or > 50 commercial antibodies; T_dark_ refers to proteins that have been manually curated in UniProt, but do not meet criteria for the above categories.

## 3. Results

### 3.1. Integrative Risk Gene Selector (iRIGS)

18 of the 98 genes (18.4%) exhibited high posterior probability (≥.75), indicating support from multiple genetic features including distance to SNP and regulatory information. 7 of these 18 genes (38.9%) were the not the closest protein coding gene to the corresponding SNP. This is consistent with prior work finding many HRGs are not the most proximal to the SNP (Wang et al., 2019). The HRGs that exhibited high posterior probability and were not the closest gene to the SNP were *GALNT17* (polypeptide N-acetylgalactosaminyltransferase 17), *AKAP13* (A-kinase anchoring protein 13), *PPP3CA* (protein phosphatase 3 catalytic subunit alpha), *KLF4* (Kruppel-like factor 4), *PRKCE* (protein kinase C epsilon), and *ZEB2* (zinc finder E-box binding homeobox 2).

Only 5 of these 18 genes (27.8%) overlapped with the 307 genes identified in the prior alcohol consumption GWAS. This indicates that 13 unique genes were identified with iRIGS (Table 1). The highest ranking genes for each SNP as well as all protein-coding genes considered within the 1Mb flanking region of each SNP are in the Supplementary Materials (Table S1 and Table S2, respectively).

**Table 1.** Genetic and target development/druggability level (TDL) information for the high-probability (≥.75) high-confidence risk gene (HRGs)

| SNP | HRG | Posterior probability | TDL | Nearest protein coding gene | Overlap of HRG with Liu et al. (2019) |
| --- | --- | --- | --- | --- | --- |
| rs62044525 | *CDH11* | 1 | T<sub>bio</sub> | *CDH11* | N |
| rs62250685 | *CADM2* | 1 | T<sub>bio</sub> | *CADM2* | Y |
| rs74664784 | *CADM2* | 1 | T<sub>bio</sub> | *CADM2* | Y |
| rs4916723 | *MEF2C* | 0.99 | T<sub>bio</sub> | *MEF2C* | N |
| rs9950000 | *TCF4* | 0.98 | T<sub>bio</sub> | *TCF4* | Y |
| rs10085696 | *GALNT17* | 0.98 | T<sub>bio</sub> | *AUTS2* | N |
| rs12907323 | *AKAP13* | 0.91 | T<sub>bio</sub> | *AGBL1* | N |
| rs10004020 | *FBXW7* | 0.90 | T<sub>bio</sub> | *FBXW7* | N |
| rs72859280 | *ACVR2A* | 0.88 | T<sub>chem</sub> | *ACVR2A* | N |
| rs4699791 | *PPP3CA* | 0.87 | T<sub>chem</sub> | *EMCN* | N |
| rs4842786 | *BTG1* | 0.86 | T<sub>bio</sub> | *BTG1* | N |
| rs10978550 | *KLF4* | 0.85 | T<sub>bio</sub> | *ZNF462* | N |
| rs11739827 | *TENM2* | 0.84 | T<sub>bio</sub> | *TENM2* | Y |
| rs1004787 | *PRKCE* | 0.83 | T<sub>chem</sub> | *SIX3* | N |
| rs13024996 | *ZEB2* | 0.82 | T<sub>bio</sub> | *ARHGAP15* | N |
| rs13383034 | *PRKCE* | 0.81 | T<sub>chem</sub> | *SIX3* | N |
| rs682011 | *SORL1* | 0.75 | T<sub>bio</sub> | *SORL1* | N |
| rs12088813 | *PDE4B* | 0.75 | T<sub>clin</sub> | *PDE4B* | Y |

*Note.* The 4 TDLs are described in the Materials and methods.

### 3.2. Target Development/Druggability Level (TDL)

Of the 18 genes identified by iRIGS, *PDE4B* was T_clin_, targeted by 8 approved drugs; *ACVR2A* (activin receptor type-2A), *PRKCE*, and *PPP3CA* were T_chem_ (i.e., small molecules bind to them with high potency); and the rest were T_bio_ (Table 1). Although *PDE4B* was already identified in the alcohol consumption GWAS, *ACVR2A, PRKCE*, and *PPP3CA* were all unique genes (Liu et al., 2019).

Of the 307 protein-coding genes identified in the alcohol consumption GWAS, 17 were T_clin_, targeted by 104 unique drugs (65 of which target *DRD2*), 29 were T_chem,_ 198 were T_bio_, and 63 were T_dark_. The 17 T_clin_ genes and their approved drugs are depicted in Figure 1. The TDLs for all 307 genes are provided in the Supplementary Materials (Table S3).

**Fig 1.**
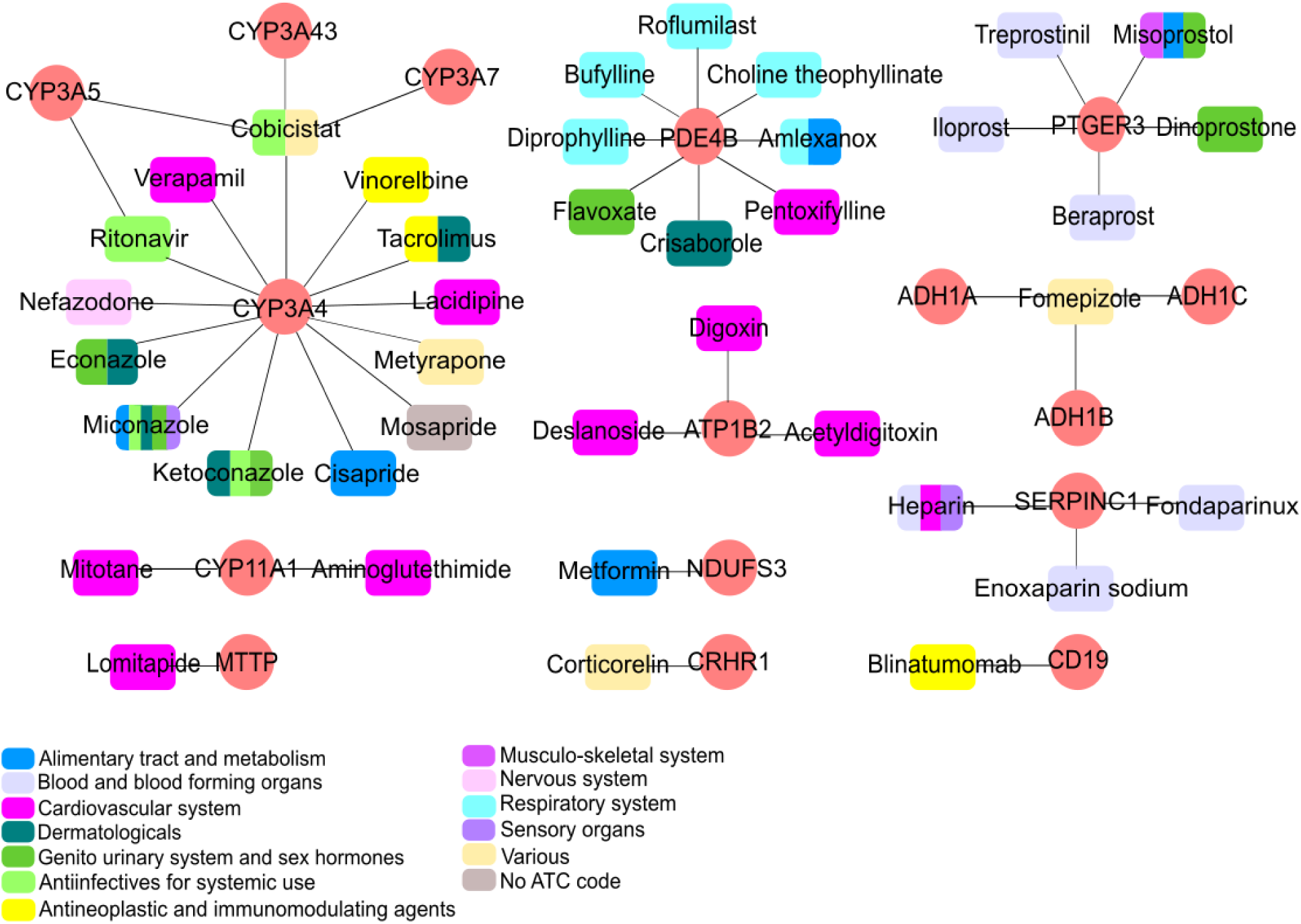
Approved drugs that interact with proteins produced by the genes identified in Liu et al. (2019). Anatomical Therapeutic Chemical (ATC) Classification System codes are color coded as indicated in the key. Medications with multiple ATC codes are assigned all of the corresponding colors (e.g., miconazole). *DRD2* is linked to 65 approved drugs (not depicted here as it is a well-established top psychiatric drug target (Liu et al., 2019; Oprea et al., 2018)).

## 4. Discussion

This study identified several novel potential risk genes for alcohol consumption and highlights putative targets for the treatment of AUD and/or ALD. In particular, *ACVR2A* codes for activin receptor type-2A, which has been linked to liver function (Haridoss et al., 2017), cocaine self-administration (Gancarz et al., 2015; Wang et al., 2017), and anxiety (Ageta et al., 2008). With regard to liver function, one study using *in vitro* models found activin A, a ligand that binds with high affinity to activin type 2 receptors, is critical to normal liver function and suggested inhibition of activin A or its downstream signaling could be a new approach for treating liver disease (Haridoss et al., 2017). Furthermore, activin A serum levels have been found to be elevated in patients with ALD compared to patients with non-alcohol related liver disease at various stages (Voumvouraki et al., 2012). Rodent studies have identified increases in activin A and activin type-2A levels in the nucleus accumbens following withdrawal from cocaine (Gancarz et al., 2015; Wang et al., 2017). Relatedly, two studies using transgenic mice expressing a dominant-negative activin receptor type-1B (also recruited in the activin A signaling pathway (Loomans and Andl, 2014)) in forebrain neurons found reduced alterations in GABAergic inhibition, hypersensitivity to the sedating effects of alcohol (Zheng et al., 2016), and low anxiety (Zheng et al., 2009).

With regard to *PRKCE*, rodent studies have found a robust link between *Prkce* expression and alcohol consumption (Ron and Barak, 2016). In particular, alcohol exposure causes changes to PKCε expression and localization in various brain regions that are implicated in addiction leading to increased alcohol tolerance and consumption (for a review see (Pakri Mohamed et al., 2018)). A recent study tested several novel molecules that act as PKCε inhibitors, finding that two promising compounds that inhibited PKCε with selectivity, crossed the blood-brain barrier, prevented alcohol-stimulated GABA release in the central amygdala, and reduced alcohol consumption in wild-type but not in *Prkce*^-/-^ mice (Blasio et al., 2018). Thus, this study provides the first human genetic support for the link between *PRKCE* and alcohol consumption, and the fact that iRIGS identified a distant gene linked to alcohol consumption that has been previously identified experimentally, supports the validity of iRIGS for identifying additional relevant genes.

*PPP3CA* codes for calcineurin, which has been found to be a regulator of GABA_A_ receptor synaptic retention and plasticity (Bannai et al., 2015; Eckel et al., 2015) and linked to diazepam response *in vitro* (Nicholson et al., 2018) and in mice (Lorenz-Guertin et al., 2019). Given GABA_A_ receptors are a primary target responsible for the effects of alcohol (Koulentaki and Kouroumalis, 2018; Roberto and Varodayan, 2017), calcineurin is likely linked with drinking via this mechanism.

This study’s analysis of the 307 genes from the prior GWAS (Liu et al., 2019) using the TDL system highlighted 17 genes that code for proteins that are targeted by at least one approved drug. Many of the drugs are promising candidates for managing liver toxicity. Fomepizole (*ADH1A, ADH1B, ADH1C*) blocks alcohol dehydrogenase and is approved to treat methanol and ethylene glycol toxicity (Ng et al., 2018). Metformin (*NDUFS3*) has been highlighted as a promising hepatoprotective agent including for ALD (Iranshahy et al., 2019). Likewise, beraprost and misoprostol (*PTGER3*), have shown promise for managing acute liver injury and liver disease (Deng et al., 2018; Gobejishvili et al., 2015; Misawa et al., 2017). Pentoxifylline has been often used to treat severe alcoholic hepatitis and a recent clinical practice update indicates that patients with a contraindication to glucocorticoids may be treated with pentoxifylline (Mitchell et al., 2017). However, this clinical practice update also notes that recent data question its clinical utility, (Singh et al., 2015; Thursz et al., 2015) thus further research is needed. Iloprost (*PTGER3*) has shown promise for managing bone marrow oedema and early stages osteonecrosis, of which excessive alcohol use is a risk factor for (Pountos and Giannoudis, 2018). Finally, PDE4 inhibitors are currently being investigated for AUD and ALD (Baillie et al., 2019; Rodriguez et al., 2019).

Numerous genes and associated drugs were also identified for the Cytochromes P450 family of enzymes, namely CYP3A4, which is implicated in the metabolism of numerous approved drugs including many antiretroviral drugs identified here (e.g., (Midde et al., 2016)). Although three small studies have examined nefazodone for AUD (Hernandez-Avila et al., 2004; Kranzler et al., 2000; Roy-Byrne, Peter P. Pages et al., 2000), interest in this medication waned given risk for severe liver toxicity in a minority of patients and an associated FDA black box warning (Edwards, 2003). Verapamil has been found to prevent cue-induced reinstatement of alcohol seeking in rats, suggesting it may be a promising compound for relapse prevention (Uhrig et al., 2017). Metyrapone has been found to prevent alcohol withdrawal associated working memory deficits and reestablish prefrontal cortex activity in withdrawn mice (Dominguez et al., 2017). In summary, the *CYP3A4* findings suggest the TCRD may be a novel method for not only identifying putative treatment targets, but also flagging potential deleterious drug-alcohol interactions.

### 4.1. Conclusions

We incorporated regulatory information and drug-protein interaction data to highlight promising molecular targets and drugs for potential repurposing. Notably, iRIGS only matched a minority of the alcohol consumption SNPs (18.4%) to genes at a confidence (≥.75) consistent with previous studies (Li et al., 2020; Wang et al., 2019), with some SNPs not matched with confidence despite being in well-replicated loci (e.g., *ADH1B*), underscoring the need to apply multiple methods to assign function to genome-wide significant loci. We focused exclusively on the SNPs from the alcohol consumption GWAS (Liu et al., 2019) in this study because it included the largest sample size and the largest number of significant loci, however, it will be important for future studies to extend these methods to GWAS of other alcohol-related phenotypes (e.g., AUD diagnosis; (Kranzler et al., 2019)) and non-European Ancestry individuals. The methods utilized within this manuscript are readily available (https://github.com/CNPsyLab/Alcohol-Genetics-iRIGS-TCRD) and applicable to other complex traits.

